# Suppression of *aedes aegypti* by sterile insect technique on captiva island, florida, usa from 2020 to 2022

**DOI:** 10.1101/2024.10.18.618919

**Authors:** Rachel Morreale, Danilo Oliveira Carvalho, Steven Stenhouse, Johanna Bajonero, Rui Pereira, Daniel A Hahn, Aaron Lloyd, David F. Hoel

## Abstract

*Aedes aegypti*, an aggressive nuisance biter and the primary vector for numerous arboviruses, such as chikungunya, dengue, and Zika, presents significant control challenges due to its ability to thrive in urban environments, escapes insecticide treatment by using cryptic resting and oviposition sites, and development of resistance to chemical mosquito control products being used routinely. To test a new approach in their integrative mosquito management toolbox. From 2020 to 2022, the Lee County Mosquito Control District (LCMCD) employed the Sterile Insect Technique (SIT) to test a new approach in its integrated mosquito management toolbox, targeting the population suppression of *Ae. aegypti* on Captiva Island, Florida. Over 24.1 million sterile males were released across three phases, covering up to 142 hectares. The study demonstrated a population reduction of up to 79% in wild adults and a 59% decline in egg densities in the primary intervention area. While population control was successful, an influx of wild females from untreated zones posed ongoing challenges to complete suppression in areas close to non-intervention areas. This supported a need for an area-wide integrated pest management (AW-IPM) approach. These results underscore SIT’s potential as a critical tool in integrated mosquito management strategies and emphasize practical application.

## INTRODUCTION

*Aedes aegypti* is a significant invasive mosquito species in warm climates across the globe because it is an aggressive biter that lives in and near human dwellings. This species can breed in a wide variety of artificial habitats and survive across a wide range of temperatures [1,2]. *Aedes aegypti* predominantly feeds on humans to complete its life cycle, and this mosquito possesses excellent vector competence and high vectorial capacity for several arboviruses, including dengue and Zika virus, among others [3,4]. *Aedes aegypti* is highly anthropophilic and is closely associated with human populations, with a strong preference for biting humans [5,6]. This close association with humans has played a vital role in the global dispersion of *Ae. aegypti*. Originating in sub-Saharan Africa and the southwestern Indian Ocean [7], this mosquito species spread overseas during European transatlantic slave trading (1500-1700), including to the American continent [8]. Historical records and descriptions of *Aedes*-borne diseases, such as yellow fever, allow us to trace the presence of the vector back in time. The first recorded instance of yellow fever in the Americas occurred in Hispaniola (present-day Dominican Republic and Haiti) in 1495, followed by a case in 1635 on Guadeloupe Island, indicating *Ae. aegypti* has long been a problem in the Caribbean [9].

Yellow fever transmission in the United States has been frequently documented since the late 1600s, with multiple seasonal epidemics in colder climate port cities like Philadelphia and Boston throughout the late 1700s and 1800s [10,11]. With the frequent transmission of yellow fever and the introduction of encephalitis over a hundred years ago, the State of Florida began a public health campaign to combat mosquito-borne disease, leading cities and counties throughout the state to form various local government organizations focused on mosquito control. The Lee County Mosquito Control District (LCMCD) was established in 1958 by merging several existing mosquito control districts from Fort Myers, Boca Grande, and Sanibel-Captiva [12]. The primary directives of LCMCD are to protect residents and visitors of Lee County, Florida, from mosquito-borne disease and nuisance biting that may detract from the enjoyment of outdoor spaces. While the incidence of mosquito-borne disease is low, Florida is a major travel destination for visitors from regions where arboviruses and malaria are highly prevalent. Each year since 2011, Florida has experienced a low incidence of local dengue transmission, primarily associated with travelers from areas where the disease is endemic dengue [13–15]. Similarly, Florida also experienced a transient outbreak of Zika with local transmission in 2016-2017 [16,17] and recently even experienced a small local cluster of malaria transmission associated with initial travel-acquired cases [18–20]. Given the high volume of international travelers visiting Florida, a sub-tropical climate, and multiple mosquito species capable of vector disease-causing agents, Florida’s mosquito control districts must be vigilant and have control strategies ready to deploy against disease-vectoring species.

The LCMCD has been employing insecticides for mosquito control since the late 1950s. Initially, the district relied heavily on DDT, a potent and long-lasting insecticide, until its environmental and health risks and mosquito resistance became well-known in the 1960s. Since then, the district has adopted various other insecticides, including malathion, naled, pyrethroids, and *Bacillus thuringiensis israelensis* (Bti), a bacterial larvicide [12]. These insecticides are applied using multiple methods, such as aerial spraying, truck-mounted spraying, and targeted applications with backpack sprayers. The district’s use of insecticides has effectively reduced mosquito populations and decreased the risk of transmission of mosquito-borne diseases. However, concerns persist regarding these chemicals’ potential environmental and human health impacts. Furthermore, resistance to commonly used larvicides and adulticides has been detected at high levels in populations of *Ae. aegypti* worldwide, including in Florida [21–23].

Given the problems associated with insecticide application and resistance development, it is crucial to explore alternative control methods. One such method currently under development is the Sterile Insect Technique (SIT) for mosquitoes, which has shown promising results based on small-scale trials in multiple countries [24–27]. The SIT involves rearing massive numbers of insect males in the mass-rearing facilities laboratory, sterilizing them, and then releasing the sterile males into the field at greater densities than wild males to increase the likelihood of females mating with sterile males rather than fertile (wild) males. With frequent and consistent releases over generations, the wild population is eventually suppressed [28].

Since 2017, in collaboration with the Insect Pest Control Subprogramme (a joint partnership between the International Atomic Energy Agency – IAEA – and the Food and Agriculture Organization – FAO), LCMCD has been developing SIT for mosquitoes. The district established baseline data collection and performed substantial public engagement on the offshore islands Captiva and Sanibel [29]. This helped define the descriptive mosquito population profile and promote acceptance of the novel intervention (publication under preparation). This is a historic place for SIT because Sanibel was where the first SIT applications against an insect pest were employed, targeting the New World Screwworm *Cochliomyia hominivorax* in the early 1950s [30,31]. To demonstrate successful SIT for *Ae. aegypti* in these historic locations, LCMCD has also performed dispersal studies and wild population estimation using the mark-release-recapture method on Captiva Island [32]. This paper focuses on the next step, demonstrating mosquito population suppression by continuously releasing sterile males and monitoring the wild population and the sterile males released [33,34]. We show that continuous releases of massive numbers of lab-reared sterile male *Ae. aegypti* can successfully suppress the wild *Ae. aegypti* population on Captiva, with suppression sustained over three years.

## RESULTS

### Site selection

The LCMCD has actively performed mosquito surveillance and control over Lee County (Figure 1A) for several decades, and Captiva Island met multiple criteria for becoming an SIT pilot area. Some of the main criteria were: 1) isolation from the mainland, reducing the chances of immigration of new *Ae. aegypti* onto the island (Figure 1B), 2) the presence of a substantial nuisance *Ae. aegypti* population with resistance to pyrethroid insecticides, 3) an *Ae. aegypti* population size [32,29] that is within our capacity to produce sufficient numbers of sterile males at the LCMCD facility to achieve population suppression, independently of the sterile:wild ratio, 4) the extremely low, sporadic, and irregular presence of *Aedes albopictus* [29], and 5) the accessibility of the area of interest for monitoring and releases of sterile mosquitoes.

**Figure 1.**
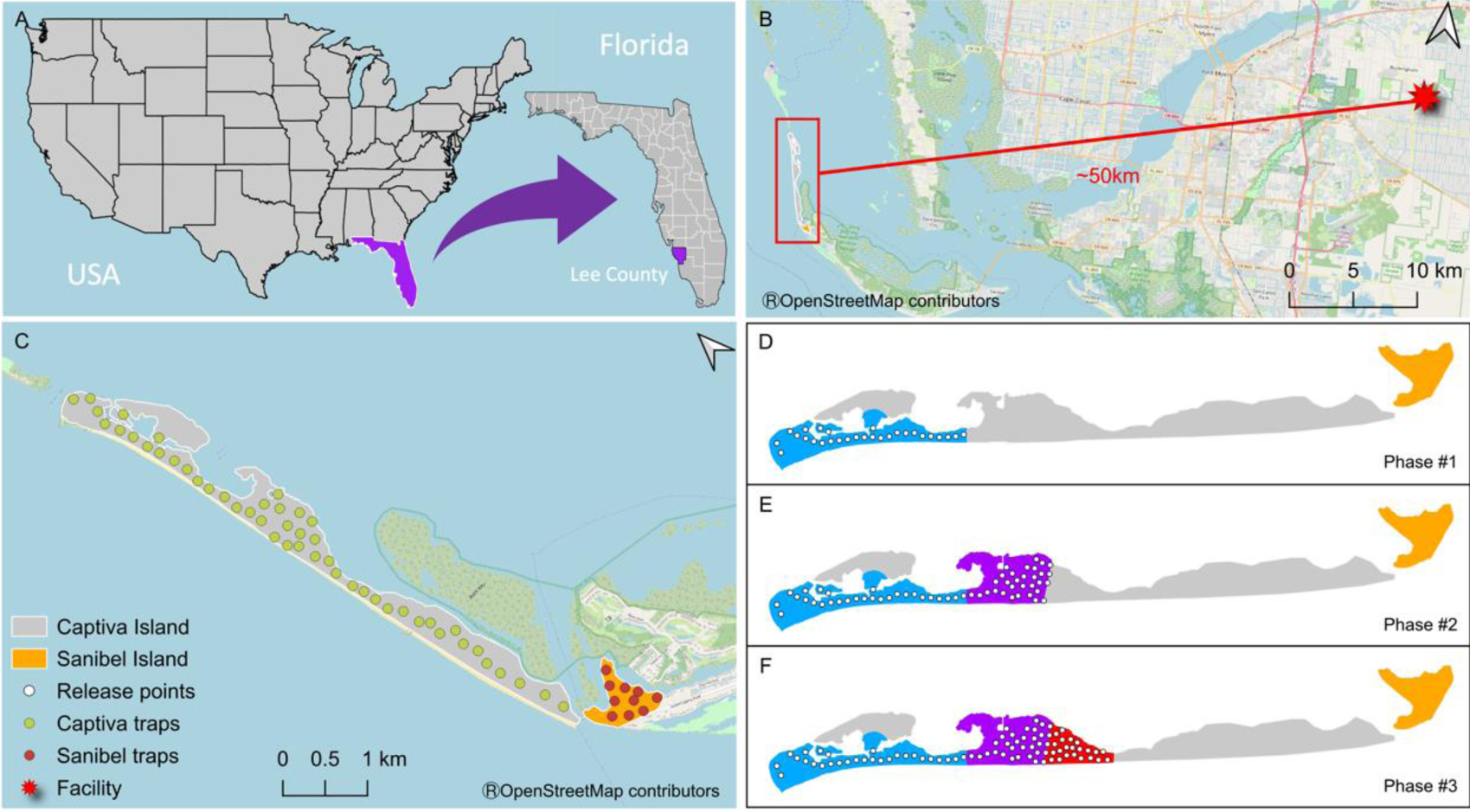
SIT study area – (A) Lee County, FL – USA. (B) The straight-line distance from the rearing facility at LCMCD and Captiva Island is around 50 km. (C) The study area on the northwestern tip of Sanibel is a non-intervention area with trapping stations, and the intervention area in Captiva has trapping stations. The intervention area was split into three release phases (D-F), showing new area aggregation and the respective release points. Trap stations (green and brown circles) comprise one ovitrap and one BGS in both intervention and non-intervention areas.

Captiva has an area of more than 278 ha (Figure 1C), with 1,108 buildings. Of those buildings, only 188 units are continuously occupied, and 920 units are transiently occupied by visitors on holiday, such that the constant human population is ∼320 year-round. However, from December through April, the island experiences high influxes of visitors at times more than tripling the number of residents on the island [35]. Because of Captiva’s large, linearly arrayed land mass and initial development of sterile male production, our release strategy was planned to start with a defined area at the island’s northern tip and increase the coverage in sequential sections over time. In this approach, we expanded the coverage area to encompass progressively more of Captiva Island as our capacity to rear more sterile males increased over time, which followed the suggestions of the phased conditional approach targeting full operational SIT implementation described previously in 2020 [34]. Similar strategies have been successfully applied for SIT programs and pilot trials for other insects [26]. Captiva Island had an intervention period from 10 June 2020 until 30 September 2022 that was divided into three phases (Figure 1D), according to the rearing capacity and expansion of the area of our releases.

### Trapping and monitoring

Our surveillance on Captiva had 48 trapping stations, each containing one BG-Sentinel 2 trap (BGS, Biogents, Regensburg, Germany) for adult collection (twice a week) and one ovitrap (OVT) for egg collection (once a week); see Figure 1C for trapping station locations. To evaluate the efficacy of our sterile male releases as a population suppression intervention, we selected a non-intervention site on the part of the neighboring Sanibel Island, connecting Captiva to the mainland through a bridge. Our non-intervention site on Sanibel is a 37.2 ha area with a similar grid of trap stations, using 10 BGS and 10 OVT, with BGS collected twice a week and OVT collected once weekly. Baseline data collection for both areas was initiated in mid-2017 to build a historical *Ae. aegypti* population profile from the islands [29]. When sterile male releases started in June 2020, the entomological surveillance efforts for egg collection for fertility assessment were increased, with ten additional OVTs in each area (intervention and non-intervention areas).

### Suppression of the wild *Aedes aegypti* population on Captiva

More than 24.1 million sterile-marked males were released in the intervention area between June 10, 2020, and September 22, 2022, a period that was divided into three phases (Phases #1, #2, and #3), as shown in Figure 1D, 1E and 1F. From the released males, 10,587 marked males were recaptured, representing around 0.02% of the total recapture rate. Releases usually occurred on Tuesdays and Fridays, according to the release schedule of the sterile-marked males. As our capacity for rearing increased, we gradually expanded the area for releasing sterile males. Table 1 illustrates the growth in our releases, highlighting the rising average number of sterile-marked males released over time, along with the expansion of the release area in each new phase.

**Table 1.** The total number of wild males and females collected in BGS fluctuates over the years. In parallel, the number of sterile males increased according to the phase requirements, wild population presence (top), and total number of eggs collected. It also shows the total number of released sterile-marked males each year. Dashes represent time points that did not exist. Before 2020, the data was considered baseline and is included in a separate publication [29].

| Activity | Area / Phase |  | 2020 | 2021 | 2022 |
| --- | --- | --- | --- | --- | --- |
| BG-Sentinel<br>Wild<br>(♂ / ♀) | Captiva | #1 | 1 361 / 2 522 | 48 / 156 | 74 / 260 |
|  |  | #2 | 827 / 1 602 | 458 / 959 | 105 / 275 |
|  |  | #3 | 3 146 / 5 579 | 3 117 / 4 909 | 1 451 / 2 194 |
|  | Sanibel |  | 4 256 / 6 688 | 5 843 / 7 751 | 1 930 / 3 275 |
| BG-Sentinel<br>Sterile males | Captiva | #1 | 1 322 | 7 679 | 2 826 |
|  |  | #2 | 25 | 1 374 | 1 983 |
|  |  | #3 | 3 | 144 | 5 589 |
|  | Sanibel |  | 0 | 8 | 12 |
| Ovitrap | Captiva | #1 | 13 350 | 2 765 | 1 077 |
|  |  | #2 | - | 49 595 | 29 936 |
|  |  | #3 | - | 79 050 | 109 464 |
|  | Sanibel |  | 100 874 | 61 801 | 206 070 |
| Sterile male<br>release | Captiva |  | 3 096 060 | 10 675 818 | 10 352 661 |

Phase #1 lasted 120 weeks with the release of sterile-marked males in our initial 63 ha release zone at the northern tip of Captiva (Phase #1), the number of wild adults captured and the number of eggs laid declined noticeably within a few weeks of our release, starting in June 2020 (Figure 2). Both wild adults captured and the number of eggs per trap remained noticeably lower than in prior years at the same site or in our non-intervention site on nearby Sanibel Island throughout our releases in 2021 and 2022 (Figure 2). The number of eggs/trap (Figure 2A) was significantly lower in our Phase #1 intervention area on Captiva Island compared to the non-intervention area on Sanibel Island. This effect was particularly apparent during the high-rain season when mosquito abundances were more significant than the low-rain season (GLM: Site (Intervention vs. Non-Intervention) z value: -37.22, ß= -2.01, and p < 0.001, Month of Suppression Trial z value: -5.21, ß= -0.01, and p < 0.001, S1 Fig A). Yet, the percentage of eggs hatching in our Phase #1 intervention area was not different between our intervention and non-intervention area (Figure 2B, GLM z value: -0.159, ß=-0.07, AND p = 0.874), suggesting either some portion of females were still mated by wild males within the intervention area or that a small number of wild females migrated in from the areas where we had not been releasing sterile males (from adjacent Phase #2). Perhaps most importantly, from a mosquito control perspective, our sterile male releases effectively suppressed the wild *Ae. aegypti* adult population (Fig 2C). When we normalized the number of wild *Ae. aegypti* captured against the number of days that BG sentinel traps were deployed in the field, defined as adult/trap/day (ATD) there were clearly fewer wild mosquitoes in our intervention zone than the non-intervention zone (GLM: Site (Intervention vs. Non-Intervention) z value: -559.6, ß= -1.87, and p < 0.001, Month of Suppression Trial z value: -716.9, ß= -0.058, and p < 0.001, Site x Month z value: -298.4, -0.081, p<0.001, S1 Fig. B). Noting the significant interaction between site and month in our model, there was also a strong effect of seasonality, with suppression of adult numbers being most apparent in the rainy season when mosquito populations are high and less apparent in the dry season when populations are naturally low (Fig 2 and S1 Fig B).

**Figure 2.**
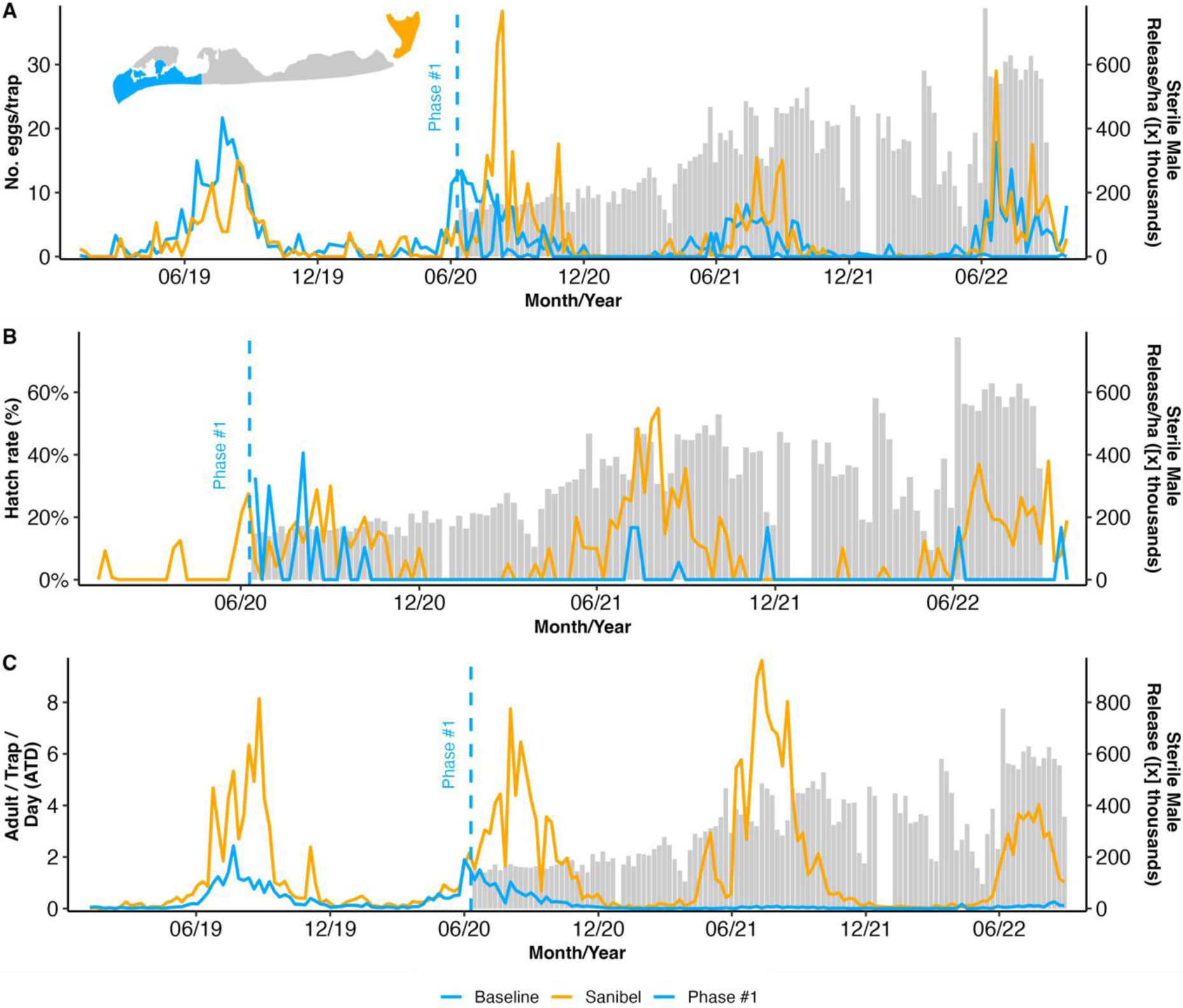
Captiva and Sanibel Islands sterile insect technique pilot (SIT) project field collection data showing the mean number of **Aedes aegypti** sterile marked males released per hectare twice a week (grey bars) and (A) the mean number of collected eggs during Phase #1 in Captiva and the non-intervention area in Sanibel Island; (B) the hatch rate of these eggs; (C) the number of adults (males and females) found per trap per day (ATD). The dashed line indicates the start of the phase’s releases.

To further test for suppression, we employed a causal impact analysis for time-series data that explicitly considers temporal autocorrelation [36]. We used adult trap captures and egg collections from the Phase #1 traps collected in the three years before releases beginning in June 2020 [29] to project a distribution of expected adult trap captures and egg collections based on these high-resolution historical data and then used the Bayesian estimation method to calculate the probability that our monitoring data after release is different from the expected population based on historical monitoring at the exact location [36,37]. We show that within weeks of our releases starting, there was a significant decrease in both the numbers of eggs captured in ovitraps and the numbers of wild adults captured in BGS traps and that the impact of releases was continued over the multiple seasons of releases (S2 Fig). The number of eggs in our ovitraps was reduced by -59% (-42 to -70% credible interval for reduction), and the effect of the intervention on eggs was -3.58 (-1.65 to -5.35 credible interval). Given these parameters for the egg population reduction, the probability of obtaining this effect by chance alone is minimal (Bayesian probability, p=0.001). Similarly, the number of adults in our BGS traps was reduced by -79% (-69 to -85% credible interval for reduction), and the effect of the intervention on the adult population was -0.53 (-0.30 to -0.77 credible interval, Bayesian probability, p=0.001).

Starting on 22 May 22, 2021, we expanded the area of intervention from the initial 63 ha of the Phase #1 area to add 54.7 ha of Phase #2 for a total of 117.7 ha intervention area with the release of sterile males over 71 weeks until the study ended (Fig 1). The expansion area for Phase #2 included the most populated area of the island. In this area, we had previously performed mark-release recapture studies [32]. Although the effects of sterile male releases on the number of eggs collected in ovitraps in Phase #2 was not as apparent as in Phase #1, there was a significant decrease in the number of eggs/trap in our Phase #2 intervention area compared to our non-intervention area (GLM: Site (Intervention vs. Non-Intervention) z value: -28.58, ß= -0.83, and p < 0.001, S1 Fig C). We determined hatching from the eggs collected in Phase #2, and similar to Phase #1, there was no difference in the proportion of eggs hatching between the intervention and non-intervention areas (Figure 3B, GLM z value: 0.322, ß=0.27, and p = 0.400). However, the number of wild adults collected in the intervention site was substantially less than in the non-intervention site in Phase #2 after releases started (Figure 3C, S1 Fig. D, GLM: Site z value: -369.7, ß= -1.29, and p < 0.001, Month of Suppression Trial z value: -716.9, ß= -0.058, and p < 0.001, Site x Month z value: -182.1, -0.044, p<0.001). Our traditional statistical approach of comparing monthly totals between our Phase #2 intervention site and our non-intervention site on Sanibel Island showed a detectable decrease in both numbers of eggs in ovitraps and, most importantly, adults captured in BGS traps (S1 Figs C and D). However, analyzing these data with causal impact analysis comparing the multi-year population census data in our Phase #2 intervention site before and after the intervention showed no detectable effect of our intervention in Phase #2 (reduction in eggs -30% but with a credible interval of -60% to +47%, the impact of -1.31 but with a credible interval of -3.15 to +0.65, p=0.094, estimates also show a reduction of -6% in adults but with a credible interval of -36% to +86% an effect of -0.01 with a credible interval of -0.16 to +0.14 and p=0.463, S3 Fig.).

**Figure 3.**
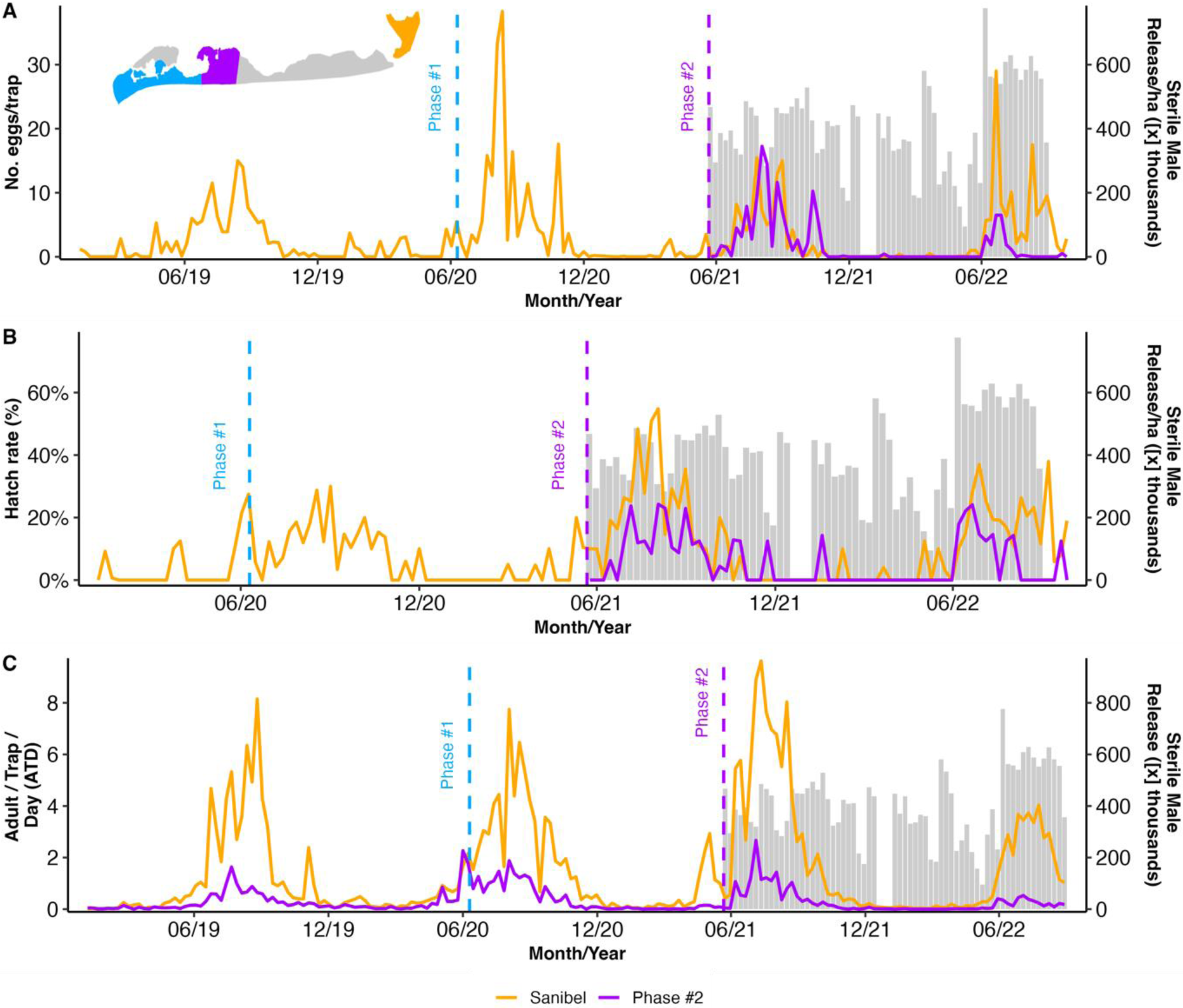
Captiva and Sanibel Islands sterile insect technique pilot (SIT) project field collection data showing the mean number of **Aedes aegypti** sterile marked males released per hectares twice a week (grey bars) and (A) the mean number of collected eggs during Phase #2 in Captiva and the non-intervention area in Sanibel Island; (B) the hatch rate of these eggs; (C) the number of adults (males and females) found per trap per day (ATD). The dashed line indicates the start of the phases’ releases.

The expansion to Phase #3 started on December 21, 2021, increasing our total area covered in sterile males to 142.2 ha (24.5 ha added) with the release of 7.1 million sterile males, and lasted 42 weeks until the study was interrupted on 26 September 2022, before Hurricane Ian hit the islands two days later. All activities ended (Figs 4A, 4B, and 4C). Over the partial season of sterile male releases, we found fewer eggs collected in ovitraps in our Phase #3 area on Captiva Island than in our non-intervention site on Sanibel Island (GLM: Site (Intervention vs. Non-Intervention) z value: -28.58, ß=-0.827, and p <0.01, Month of Suppression Trial z value: - 22.4, ß=-0.0241, p<0.001, S1 Fig E), no effect on egg hatching (GLM: Site z value: 1.14, ß=0.26, and p = 0.252), and a small but detectable effect on adults collected in BGS traps (GLM: Site z value: -292.9, ß= -0.64, and p < 0.001, Month of Suppression Trial z value: - 716.9, ß= -0.058, and p < 0.001, Site x Month z value: -2.84, ß= -0.004, p=0.0045, S1 Fig F). As for Phase #2 above, our traditional statistical approach of comparing monthly totals between our Phase #3 intervention site and our non-intervention site on Sanibel Island showed a detectable decrease in both numbers of eggs in ovitraps and adults captured in BGS traps but with admittedly small effect sizes. Causal Impact Analysis (CIA) comparing our Phase #3 intervention site before and after the intervention showed no detectable effect of our intervention in Phase #3 (reduction in eggs of -32% but with a credible interval of -66% to +69%, effect of -1.32 but with a credible interval of -3.24 to +0.70, p=0.09, estimates show a reduction of -19% in adults but with a credible interval of -45% to +26% an effect of -0.19 with a credible interval of -0.52 to +0.13 and p=0.139, S4 Fig).

**Figure 4.**
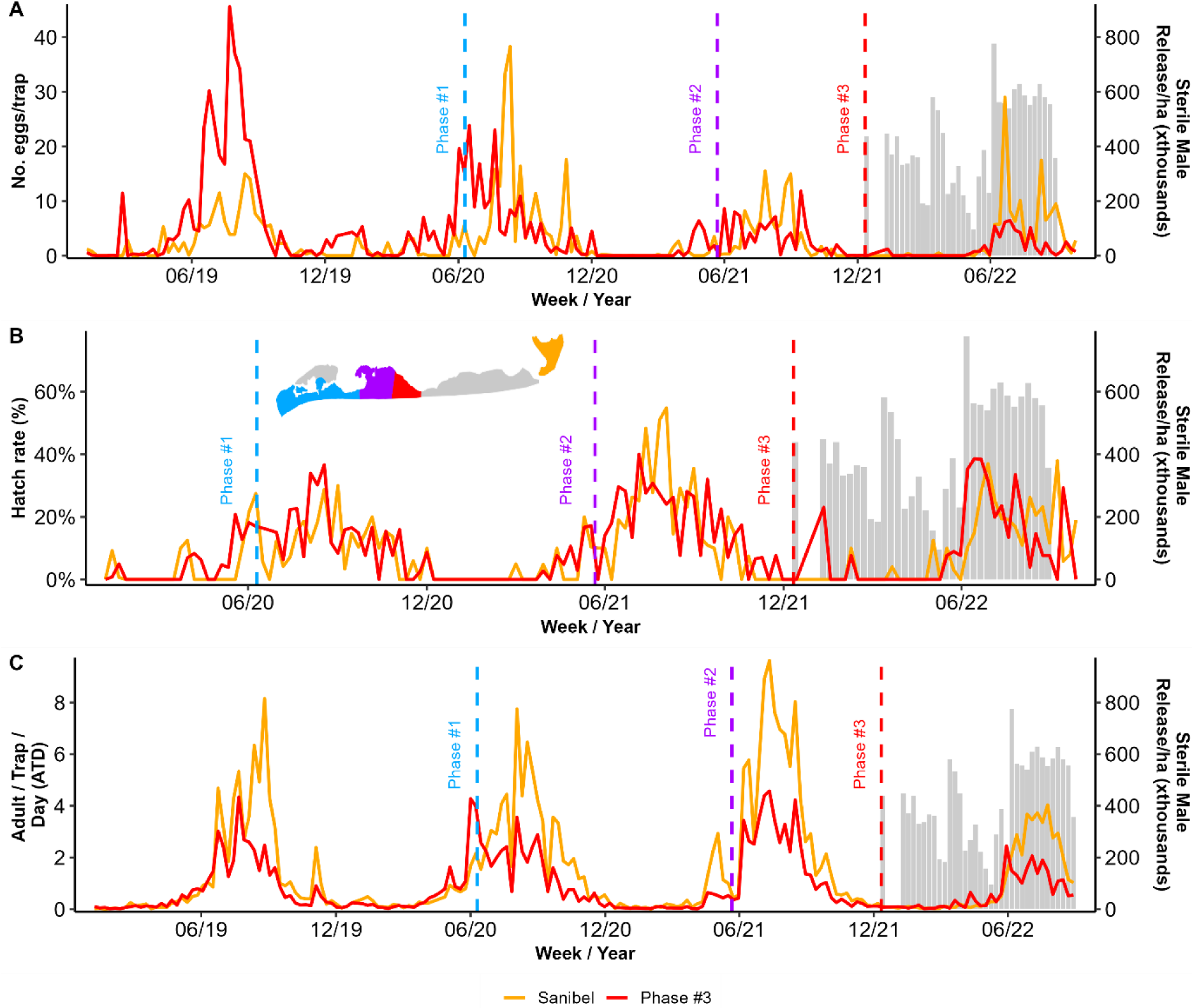
Captiva and Sanibel Islands sterile insect technique pilot (SIT) project field collection data showing the mean number of **Aedes aegypti** sterile marked males released per hectares twice a week (grey bars) and (A) the mean number of collected eggs during Phase #3 in Captiva and the non-intervention area in Sanibel Island; (B) the hatch rate of these eggs; (C) the number of adults (males and females) found per trap per day (ATD). The dashed line indicates the start of the releases of the phases.

### Ratios of sterile to wild males

During the three phases of our suppression trial, we released more than 24.1 million sterile males on Captiva overall. From the total released, 0.02% of the adults were recaptured (10,587 marked sterile recaptured adults). We defined the sterile-to-wild male ratio as the number of recaptured marked sterile males collected for each wild male. Across the release phases, the mean ratio typically ranged from 0.81 sterile:1 wild up to 13 sterile:1 wild, although it occasionally reached higher values (104.7 sterile:1 wild ratio). The ratio of sterile to wild males was relatively high during the dry season when wild populations are naturally low on Captiva [29]. The ratios of sterile to wild males were relatively low at times of the year when wild populations were high in the wet seasons of each year. Yet, we were able to affect population suppression in phase #1 despite these low sterile:wild ratios (Fig 5, Table 2).

**Figure 5.**
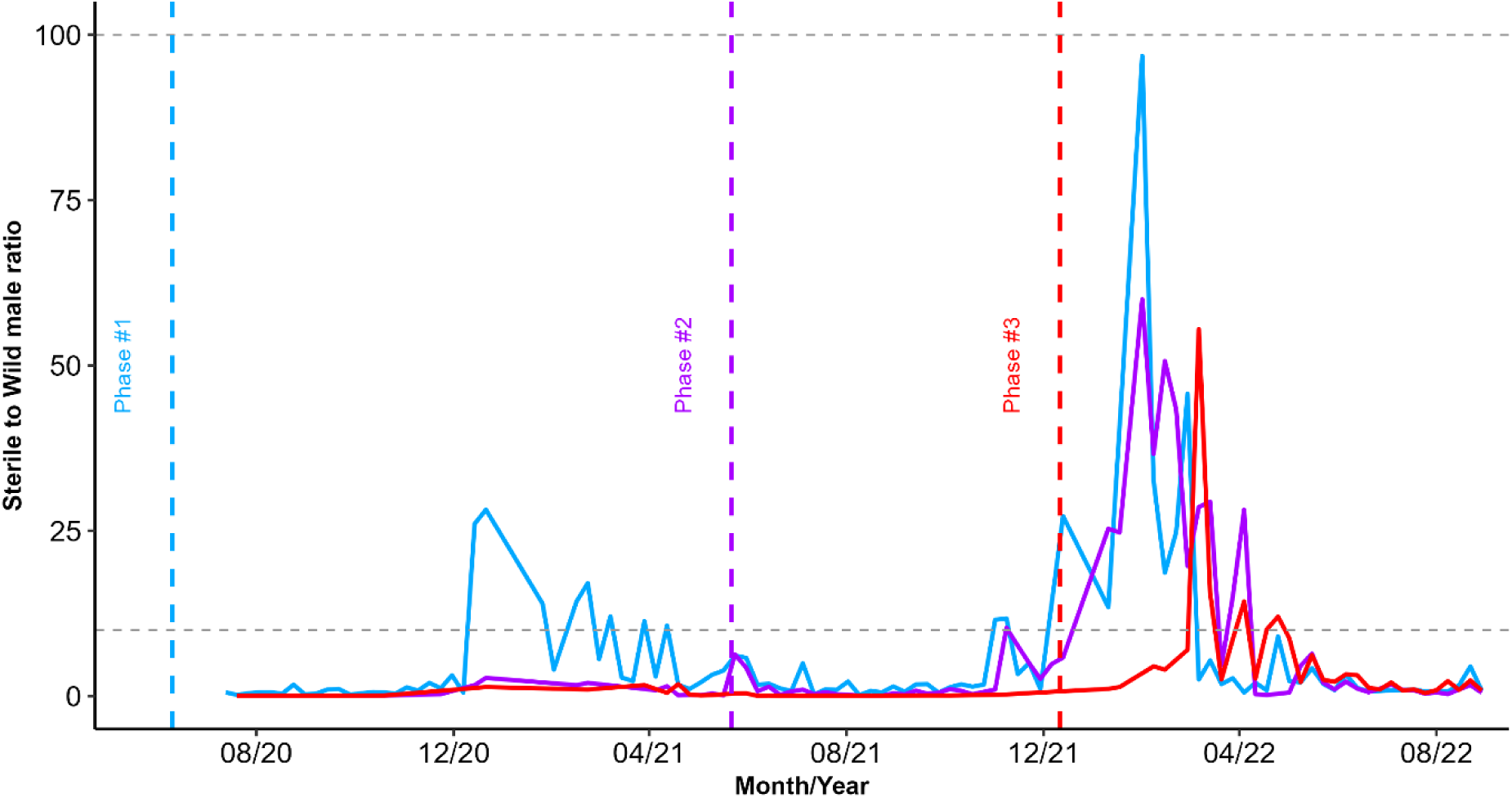
Seasonal male ratio—the proportion of sterile males to one wild male recaptured from BG-S between 2020 and 2021. Dashed lines represent the literature-recommended interval for SIT to reach successful suppression.

**Table 2.**
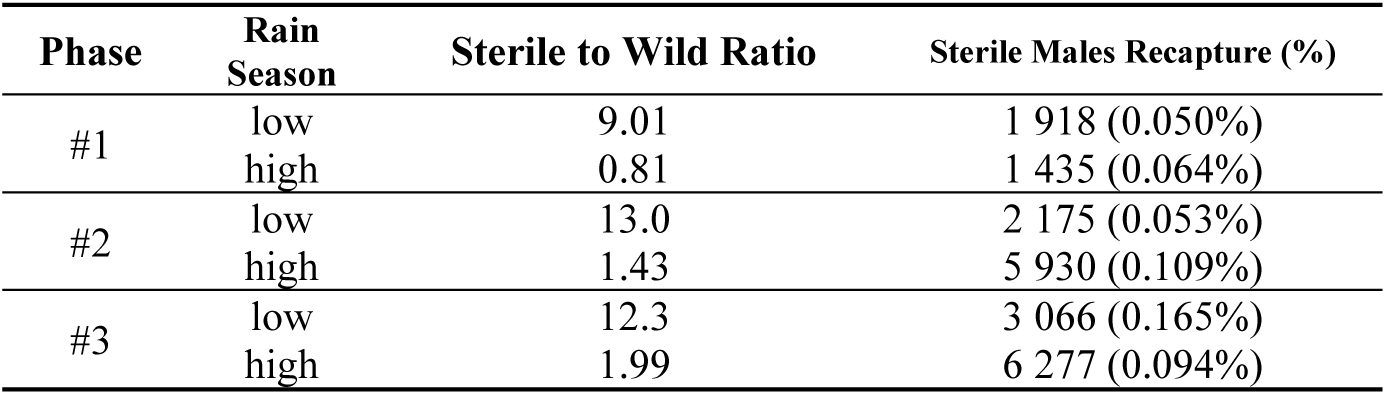
Seasonal mean sterile to wild male ratio and recapture – the proportion of sterile males to one wild male recaptured in BGS from 2020 to 2022 with the respective recapture rate.

| Phase | Rain Season | Sterile to Wild Ratio | Sterile Males Recapture (%) |
| --- | --- | --- | --- |
| #1 | low | 9.01 | 1 918 (0.050%) |
|  | high | 0.81 | 1 435 (0.064%) |
| #2 | low | 13.0 | 2 175 (0.053%) |
|  | high | 1.43 | 5 930 (0.109%) |
| #3 | low | 12.3 | 3 066 (0.165%) |
|  | high | 1.99 | 6 277 (0.094%) |

## DISCUSSION

Our initial long-term goal for this project was to suppress *Ae. aegypti* across the entirety of Captiva Island. Captiva is a relatively long and narrow barrier island that runs roughly north to south along the west coast of Florida, as seen in the map in Figure 1. Because our facility could not initially produce enough sterile males to cover the entirety of Captiva Island, we chose to break the island into sectors or phases by which we could expand the coverage of the area of releases as our rearing capacity increased as well, as has been suggested by several best practices documents for the implementation of mosquito SIT programs [34,38]. We began with our first phase at the northern end of the island and initially covered 63 ha starting in June 2020, with additional area covered in two subsequent phases. Our second phase began in May 2021 once our rearing capacity increased, expanding our sterile male coverage area to 54.7 ha. Then, our third phase started in December 2021 and expanded our area of releases to an additional 63 ha, with a total release area of 142.3 ha, or roughly 51.2% of the 278 ha island. Unfortunately, due to the catastrophic damage to the island by Hurricane Ian in September 2022, we decided to end our program before the entirety of the island was covered by sterile male releases. Thus, our results must be interpreted in the context of each of the three phases of releases, specifically because Phase #1 lasted over three full seasons, Phase #2 lasted only two seasons, and Phase #3 less than one entire season before the project was terminated (Fig 4). Furthermore, the spatial structuring of the phases was such that Phases #2 & #3 acted as buffer zones for Phase 1, separating the area of Phase #1 from the remainder of the island that received no sterile male releases.

We were able to successfully suppress the population of *Ae. aegypti* in our Phase #1 field site on Captiva Island within a few months of beginning sterile male releases, and we maintained excellent control for a total of three mosquito active seasons before ending our program due to the catastrophic effects of Hurricane Ian (June 2020 - September 2022). Mosquito abundance on Captiva is highly seasonal, with very few adults captured during the dry season, typically from late October to early May, and populations proliferating rapidly and showing high activity levels during the rainy season, generally mid-May to early October [29,32]. Perhaps most importantly, we showed a dramatic decrease in adult mosquitoes in BGS traps in our Phase #1 release area over the first season of high mosquito activity with almost undetectable *Ae. aegypti* adult populations in the second and third release seasons (Fig 2C). In the case of our Phase #1 area, we showed substantial declines in the numbers of wild mosquitoes captured in BGS traps both when comparing the intervention zone on Captiva to our non-intervention zone on nearby Sanibel Island and also by using causal impact analysis to leverage our three years of prior monitoring of these same traps [29]. Thus, monitoring mosquito populations in our core location for three years before beginning sterile male releases gave us excellent precision to show a significant suppression effect with high confidence. From a mosquito control perspective, by substantially decreasing the number of adults in traps, we expect that we effectively suppressed biting pressure from *Ae. aegypti* in our Phase #1 site on Captiva using sterile male releases.

Similarly to our successful suppression of adult numbers in our Phase #1 site, our sterile male releases decreased the number of eggs collected in ovitraps over the first season of releases. That suppression of the number of eggs collected from ovitraps was further maintained in the second and third seasons of field releases (Fig 2A). Despite our successes in reducing the number of eggs in ovitraps and the numbers of adults in BGS traps, we could not detect a reduction in the numbers of eggs hatching in our Phase #1 intervention site during our three-year trial. It is possible that the few females remaining in the Phase #1 intervention site were successfully mating with wild males rather than sterile males. However, we think it is most likely that the fertile eggs found in our Phase #1 intervention site were laid by females that had migrated into the Phase #1 area from the areas just south of our release zone where sterile male releases had not yet started, despite the buffering of our release areas for Phases #2 & #3. Phase #2 was 54.7 ha, and Phase #3 was 24.5 ha, but the linear distance between the edge of Phase #1 and Phase #2 was only 1.0 km, and Phase #3 was only 0.86 km. Although, several reports show that *Ae. aegypti* adults are not particularly strong fliers [39–41]; our own mark-release-recapture study on Captiva showed a mean traveled distance of around 201.7 m [30], and over several days, females could have dispersed from our non-intervention area into our Phase #1 intervention zone by combining flight and wind dispersal. Furthermore, human activities may also assist *Ae. aegypti* movement within Captiva, such as the natural flow of vehicular traffic from outside our release zone, as cars tend to move towards the high concentration of restaurants in our Phase #2 area of Captiva. Additionally, the resort area in Phase #1 had a shuttle that could have efficiently transported mosquitoes into and out of the suppression area as it ferried tourists between stops in the non-intervention zone and returned to the intervention zone.

Although the suppression was not as strong as in our Phase #1 area, in our Phase #2 area, we did observe a statistically detectable decrease in both the numbers of adult *Ae. aegypti* captured in BGS traps and the numbers of eggs collected in ovitraps (Fig 4A, Fig 4C, and S1 Fig). The reduction in unmarked, wild adults per BGS trap was apparent within the first season of releases in our Phase #2 area compared to our non-intervention area on Sanibel. Still, the reduction in the number of eggs per ovitrap was not apparent until releases continued into the second wet season with high adult numbers. We believe that our Phase #2 release area may have helped promote more effective suppression in our Phase #1 area by acting as a buffer zone, reducing the immigration of wild female adults into our Phase #1 zone. However, we did not detect any statistically detectable decrease in egg hatching in our Phase #2 zone, likely due to the immigration of wild females from areas of Captiva where releases were not occurring. Given that the total number of eggs in our Phase #2 intervention area was reduced substantially in the second season of releases, immigration of a relatively few fully fertile wild females into an intervention zone can provide enough fertile eggs to prevent any statistically detectable effect on the proportion of eggs hatched [42,43].

Our releases in Phase #3 began before the mosquito active season. They terminated approximately 80% of the way through the active, wet season in late September 2022 with the catastrophic landfall of Hurricane Ian on Sanibel and Captive Islands, as well as much of southwestern Florida. Similarly to our observations in our Phase #2 area, we observed a statistically detectable reduction in the number of adults caught in BGS traps within the release period in our Phase #3 area compared to our non-intervention zone on Sanibel Island (S1 Fig). However, we did not observe a detectable decrease in the number of eggs collected in ovitraps in our Phase #3 zone, likely due to the immigration of wild, fertile females from the areas south of our Phase #3 zone on Captiva where sterile-male releases had not yet begun. As mentioned above, we expect that our Phase #3 release area would act as a buffer zone for Phases #2 and #1 in summer 2022, facilitating the decrease in eggs laid in ovitraps in our Phase #2 zone in the active season when our Phase #3 releases were also occurring. Buffer zones, wherein sterile males are released in an area surrounding a core zone for interventional population suppression, have been used in several other studies of both radiation-sterilized male mosquitoes and *Wolbachia*-infected incompatible male mosquitoes and help increase the efficacy of suppression in the core intervention zone [38,44–46]. Furthermore, buffer zones for SIT interventions can consist of sterile males released in areas adjacent to a focal intervention area or other strategies for integrated mosquito management, such as larvicide or adulticide applications [47].

Despite achieving significant suppression of the *Ae. aegypti* population, our Phase #1 study operated with a notably low sterile:wild ratio. This outcome challenges the conventional recommendation ratio between 10:1 and 100:1. Our results suggest that other factors may have contributed to the success of the SIT program. The quality and competitiveness of the released sterile males were likely high, easing effective mating with wild females and resulting in population suppression. Additionally, specific environmental conditions and the dynamics of the local mosquito population may have enhanced the efficacy of the sterile males [48].

While the recommended sterile:wild ratio aims to ensure a high probability of sterile males mating with wild females, our findings indicate that lower ratios can still be effective under certain conditions. This insight could be valuable for future SIT programs, suggesting that with optimal release strategies and high-quality sterile males, population suppression can be achieved with fewer resources. Further research and analysis should focus on determining specific conditions and methods that could refine SIT guidelines and improve the efficiency of mosquito control efforts globally [49].

Many other authors have pointed out that SIT is not a stand-alone treatment [44,50]. Instead, it is crucial to conceive of SIT as one tool in the more extensive toolbox of integrated mosquito management. Most mosquito control organizations use relatively broad-spectrum chemical pesticides, either adulticides or larvicides, that can suppress multiple target species based on what species are most prevalent at any particular time of the year [51–53] thus, species-specific techniques like SIT should be reserved for control of challenging species, such as *Ae. aegypti*, that tends to be difficult to control with typical chemical applications. These difficulties lie with their choice of cryptic resting locations often protected from ultra-low volume adulticide sprays, the scattered nature of larval breeding sites that are frequently difficult to target with larvicidal treatments, and the evolution of resistance to common chemical pesticides [54,55]. SIT can be combined with other mosquito-control tools to create one solid integrated mosquito management program. For example, beyond *Ae. aegypti*, the saltmarsh mosquito *Aedes taeniorhynchus* is a critical nuisance pest on Captiva and Sanibel Islands, as well as in other coastal areas in Florida, due to its emergence in large numbers, ability to disperse 10 km or more from breeding sites to feed, and aggressive evening biting that can be a significant disruption to outdoor recreation and tourism in oceanside towns in Florida [56]. During the course of this SIT population suppression study, LCMCD applied larvicides and adulticides to Sanibel and Captiva Islands as needed in order to suppress populations of pestiferous mosquitoes and to prevent or limit arboviral disease transmission. As LCMCD is mandated by the state of Florida (FL Stat § 388.162 (2023)) to maintain and control mosquito populations for public health and quality of life, we could not withhold pesticide applications in the study areas. As a fully integrated mosquito management program, traditional mosquito suppression occurred alongside sterile male releases. Chemical treatments that were likely to impact populations of *Ae. aeg*ypti were recorded and are provided as supplementary information (S7 Table). Applications of larvicides that would not affect the larval habitats used by *Ae. aegypti* were not included in the table but took place in salt marsh and vegetated areas in and around the study areas.

Working as a part of an established and well-rounded mosquito control district following the best practices for integrated mosquito management allowed us to implement our sterile male releases better. The ability to schedule releases or treatments around each other provided a distinct advantage and, we believe, contributed to the overall success of the pilot study. In general, because sterile male releases occurred on a regular and scheduled basis, treatments with adulticide could be arranged to be conducted on different days than the preplanned releases of sterile males. Occasionally, we had to delay a release when urgent adulticiding was required. Still, due to prompt and clear communication between the surveillance and adulticiding departments in our district, sterile male release events rarely overlapped with adulticiding treatments. A further example of the integration of chemical control treatments with our SIT project occurred just before the beginning of releases. Due to the impacts of COVID-19, we could not begin releases when populations were seasonally lowest in spring 2020, as is generally advised [32]. However, using low-volume larvicide followed by adulticide treatments before releases in the active season allowed us to begin with lower populations when we began releases. Fully employing area-wide population suppression efforts allowed us to mitigate *Ae. aegypti*’s rising populations better when we started releasing sterile males in June 2020.

Conducting our own trapping and locally rearing mosquitoes in-house was also beneficial for our program. Adjusting the amounts of released males based on promptly collected field data allowed for responsive and prescriptive releases. Because mosquito production was local to the release area, males were not subjected to additional stressors that can be induced through shipment and prolonged transportation [36]. Having an independently operating SIT program with a dedicated staff allowed for a more robust program overall, consistently working to improve our methods and facilitating our scale-up over time, thus contributing to successful population suppression efforts with the SIT.

## MATERIAL AND METHODS

### Ethical committee

The creation, existence, and activities of the Mosquito Control Districts in Florida are regulated under the 2018 Florida Statutes, title XXIX (Public Health), Chapter 388 (Mosquito Control), in sections 171 and 181, which state the “*Power to perform work*” and the “*Power to do all things necessary*”, in which states “*The respective districts of the state are hereby fully authorized to do and perform all things necessary to carry out the intent and purposes of this law*”. With that, no further authorization was necessary to develop and apply mosquito trapping within the County. No human or vertebrate animal subjects were used in this work.

### Pilot site - Captiva and Sanibel Islands

Captiva and Sanibel Islands are in Lee County (Florida, USA—26.518028, -82.191057). Two other publications have already described the area: one describes the baseline data collection [29], and the second estimates the population parameters by mark-release-recapture studies [32]. Due in part to geographic isolation, the islands were known to have the presence of *Aedes aegypti* without the continuous presence of *Aedes albopictus*. Isolation is an essential factor for the SIT application, limiting the migration of mosquitoes in and out of the selected areas.

### Mosquito surveillance

The surveillance network was described in depth in a previous publication describing the period before releases, also known as the baseline data collection period [29]. Surveillance began on June 22, 2017, and continued until September 22, 2022. Adult surveillance occurred using forty-five BG-Sentinel 2 traps placed 200 m apart throughout Captiva, with the southernmost kilometer receiving three traps placed 300 m apart. The northwesternmost neighborhood on Sanibel served as the control area, with ten traps placed 200 m apart. Each surveillance site also had a corresponding oviposition trap maintained weekly to monitor egg levels (after egg sterility release). Once releases began, ten additional oviposition traps were added to the release area and ten to the control area to intensify egg surveillance out of concern that oviposition might be missed.

Adult mosquitoes collected from the field were identified at the species level. Ovitrap papers collected were allowed to cure over a week and then quantified for the number of hatched, viable-appearing, and collapsed eggs. Beginning August 13, 2019, through October 13, 2020, fifteen ovitrap papers were randomly selected. If eggs were present, these papers were flooded with 250 ml water, with hatched larvae reared to adults and identified to species. From October 20, 2020, through the end of the study, all ovitrap papers with eggs were evaluated using this process to better estimate the induced sterility in the field.

### Mosquito mass-rearing

A colony of *Ae. aegypti* was domesticated using eggs collected from the study areas in 2017, with the introduction of wild *Ae. aegypti* from these exact locations every ten generations as part of the colony management to maintain high-quality standards. Mosquitoes were reared and maintained at 26.7 ± 3°C with 80 ± 2% RH and a 14:10 L:D photoperiod. Adult colonies were provided with 10% sucrose *ad libitum,* and females were allowed to feed blood using methods similar to those described by others in our organization [57]. Eggs were hatched using a nutrient broth composed of 100 mg fish fry powder (AquaMax Fry Starter, Purina Animal Nutrition, Arden Hills, MN, USA) and 100 mg brewer’s yeast in 500 ml water, which had been incubated for four hours [58]. Approximately 16 h after flooding the eggs, the larvae and hatch solution were transferred into mass-rearing racks (Wolbaki Biotech Corporation, Guangzhou, Guangdong Province, China). Larvae were reared at 2.3 larvae/ml density and fed a slurry of fish fry powder daily at increasing rates, as described in a previous publication [32]. Pupae were drained and separated using a glass plate separator (John W. Hock Company, Model 5412, Gainesville, FL, USA). After October 23, 2020, separations were made using a Wolbaki automated pupae sex sorter (Wolbaki Biotech Corporation, Guangzhou, Guangdong Province, China). Subsamples of pupae were visually inspected to ensure a female contamination rate of 0.2% or less.

### Irradiation

As described previously, male pupae were prepared for irradiation by drying them on paper in a food dehydrator to remove excess water [59]. Dried pupae were aliquoted into groups of 2200 individuals and placed in 3 x 8 cm polystyrene tissue culture plate lids (CELLTREAT Scientific Products, Ayer, MA, USA) modified with fiberglass screen in the center for irradiation. Pupae were irradiated at 42 Gy using an X-ray irradiator (RS-2400, Rad Source Technologies, Buford, GA, USA). The absorbed dose was measured using Gafchromic MD-V3 film dosimeters (1 x 1 cm) (Ashland, Covington, KY, USA, uncertainty below 2%), following the guideline from IPC-FAO/IAEA [60]. The films were read ∼24 h after irradiation using a DoseReader 4 spectrophotometer (Radiation General Instrument Development and Production Ltd., Budapest, Hungary).

After irradiation, the aliquots of male pupae were placed in 2.37 L cylindrical plastic emergence containers prefilled with 200 ml of water. Lids, modified with a screen in the center, were placed on the containers. After emergence, adult males were provided 10% sucrose solution *ad libitum* until release. Once emergence was complete, the water was drained through a small slit cut in the side of the container and retained to assess mortality on any remaining pupae or failed emergence. Emergence containers were allowed to dry completely (∼16 h) before marking.

## Marking and release strategy

### Sterile male marking

Passive marking through emergence in fluorescent powder-conditioned-coated pots (Techno Glow, Ennis, TX, USA) was used from June 10, 2020, until July 8, 2020. Marking via insufflation was used from July 10, 2020, through the completion of releases on September 26, 2022. As of December 8, 2020, males were marked using DayGlo Eco pigments (DayGlo, Cleveland, OH, USA). Methods for both of these techniques can be found in more detail [32]. Marking methods changed to insufflation to reduce the amount of fluorescent powder used for mosquito marking, along with improved male quality and male survival as less excess powder reduced mortality.

### Marked males in Petri dishes

Releases from June 10 to June 24, 2020, had pupae emerge in fluorescent powder-conditioned emergence pots, according to methods described previously [32]. These pots were chilled in a refrigerator at ∼ one ℃ for ten minutes; then, males were dislodged and transferred to a Petri dish. A disk of paper towels was placed into the Petri dishes before the addition of mosquitoes to prevent condensation from causing them to stick to the Petri dish. Petri dishes were secured with a rubber band and put into a portable cooler (XtremePowerUS, model #94018, Chino, CA, USA) set at 11 ℃. The cooler was transferred to a vehicle (van) and driven to the release area on Captiva (around 60 minutes). Once at a release site, one Petri dish was removed from the cooler, and the lid was removed. The dish was warmed by hand, and males were encouraged to disperse by gently tapping on the bottom of the Petri dish. After ∼ 5 min, most sterile males had dispersed, and the Petri dish was placed onto a plastic plant saucer to allow any remaining live males to fly away. Vegetable oil was added around the Petri dish to deter ants and lizards from consuming the remaining males. Dishes and saucers were collected in the same order set after all release points received males. The remaining males were retained and assessed for mortality in the laboratory.

### Marked males in emergence pots

Beginning July 15, 2020, all sterile males were released directly from the emergence containers because this was faster overall. Release methods were consistent with those published previously [32].

### Releases

Before the release of sterile males, all areas received an area-wide low-volume larvicide treatment of Vectobac® WDG at 0.37 lb/acre on June 5, 2020. This was followed by an aerial ultra-low-volume Dibrom® adulticide application on June 8, 2020. These treatments were applied to reduce *Ae. aegypti* populations in preparation for the SIT releases. Releases of sterile male *Ae. aegypti* began on June 10, 2020. It was noted in previous mark release recapture (MRR) studies that sterile males had a mean distance traveled of 201.7 m and an average life expectancy of ∼2.46 days [32]. To ensure good spatial coverage of released sterile males, release points were set approximately 100 m apart.

Releases occurred twice per week, usually on Tuesdays and Fridays. Releases began at approximately 09:30 and continued until all had been released, around 13:00 (without interruptions or intervals). Approximately 100,000 sterile males/week were released initially. This amount increased as production capabilities improved over time, with over 350,000 sterile males/week towards the end of the study. Releases began at the northernmost point on Captiva and continued south throughout most of South Seas Island Resort, comprising around 15 release points. As male production increased, the release area was gradually expanded. The initial release area was ∼ 63 ha and increased to 142.2 ha when releases concluded. Males who did not fly out of the release container were retained, and mortality was quantified when they returned to the laboratory. Applications of sterile males were initially distributed as evenly as possible. Once differences in wild mosquito activity began to be apparent and the number of mosquitoes produced in the laboratory supported it, particular release points (hotspots) would receive an increase in sterile males, often either double or triple the initial release amounts, based on consistent field data from previous weeks.

### Statistical analysis

Statistical analyses were conducted using R within the RStudio environment [61,62]. The results of these analyses are provided as supplementary material (S5 Table and S6 Text). The relevant R packages and functions used are also included in the scripts in the code provided in the S8 File. Maps were created using QGIS version 3.16.9 Hannover, with a background map sourced from OpenStreetMap (Creative Commons Attribution-ShareAlike 2.0 license). Generalized linear models (GLMs) were employed to evaluate male recapture parameters, such as recapture percentage and the number of recaptured males and females, egg analysis, and hatch rate. A binomial distribution was used for percentages, while a Poisson distribution was applied for counts. The Causal Impact Analysis (CIA) was used from the R package [36,37] as the interrupted time-series analysis to determine whether releases impacted the wild population.

## ACKNOWLEDGMENT

We thank LCMCD technicians for their support in trapping, processing, and recording data. This study was supported by the Insect Pest Control Subprogramme of the Joint FAO/IAEA Centre of Nuclear Techniques in Food and Agriculture and the United States State Department in the frame of the “Surge Expansion of the Sterile Insect Technique (SIT) to Control Mosquito Populations that Transmit the Zika Virus” project.

## AUTHOR CONTRIBUTIONS

**Conceptualization:** RM, DOC, RCP, AL, DFH

**Methodology:** DOC, RM

**Software:** DOC

**Validation:** RM

**Formal analysis:** DOC, DAH, RM

**Investigation:** RM, DOC, SS, JB

**Resources:** RM, SS

**Data Curation:** RM, SS, DOC, JB

**Writing - Original Draft:** DOC, DAH, RM

**Writing - Review & Editing:** DOC, RCP, JB, DAH, AL, DFH

**Visualization:** DOC, DAH

**Supervision:** RM, RCP, AL, DFH

**Project administration:** RM, AL, DFH

**Funding acquisition:** RM, AL, DFH

## SUPP. MATERIAL

### SUPPLEMENTARY Figures

**S1 Figure.**
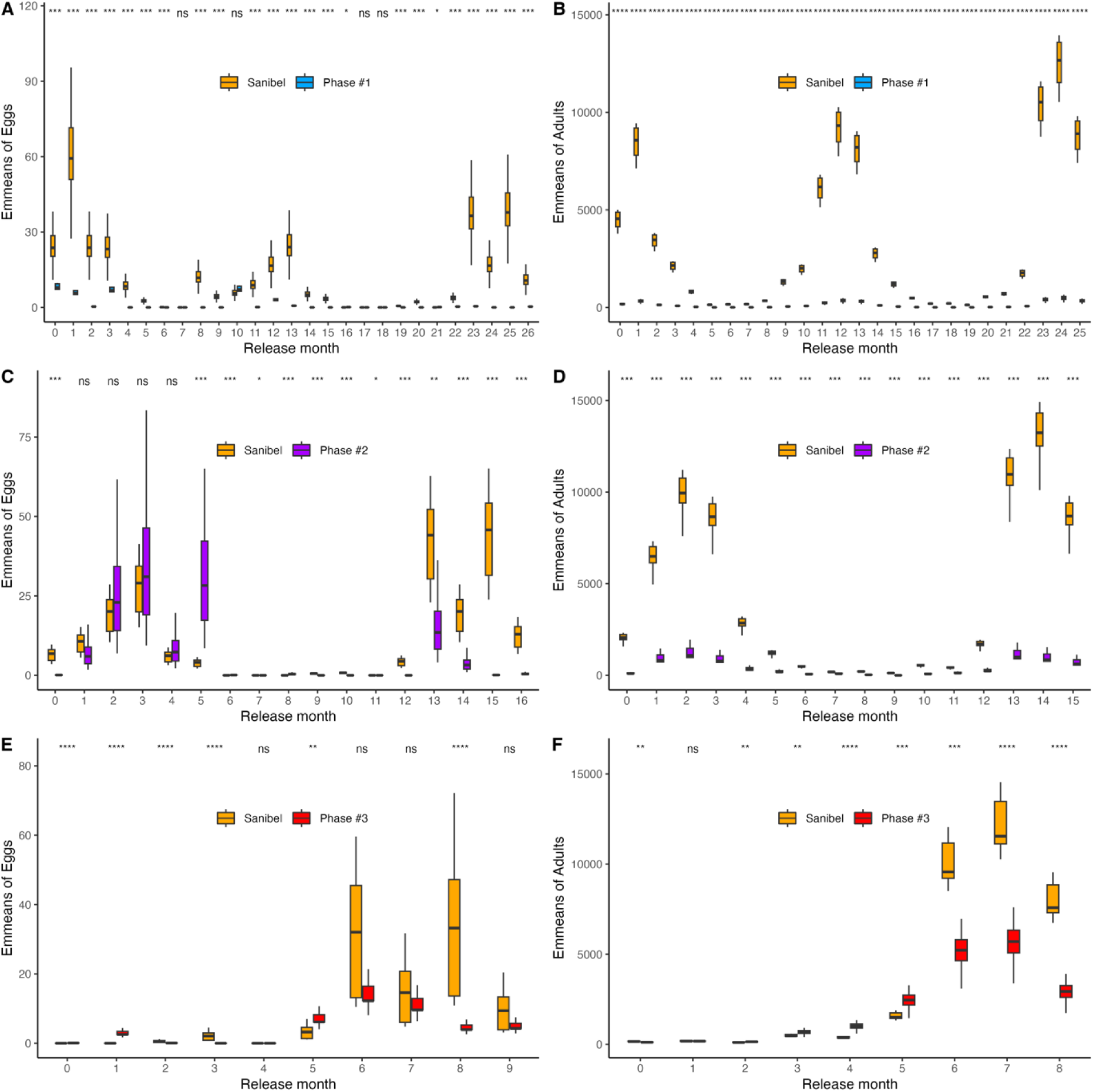
Emmeans (obtained from a zero-inflated analysis) comparing the intervention and non-intervention areas considering the beginning of the releases in each phase (1, 2, and 3) for the egg density (A, C, and E) and adult density (B, D, and F).

**S2 Figure.**
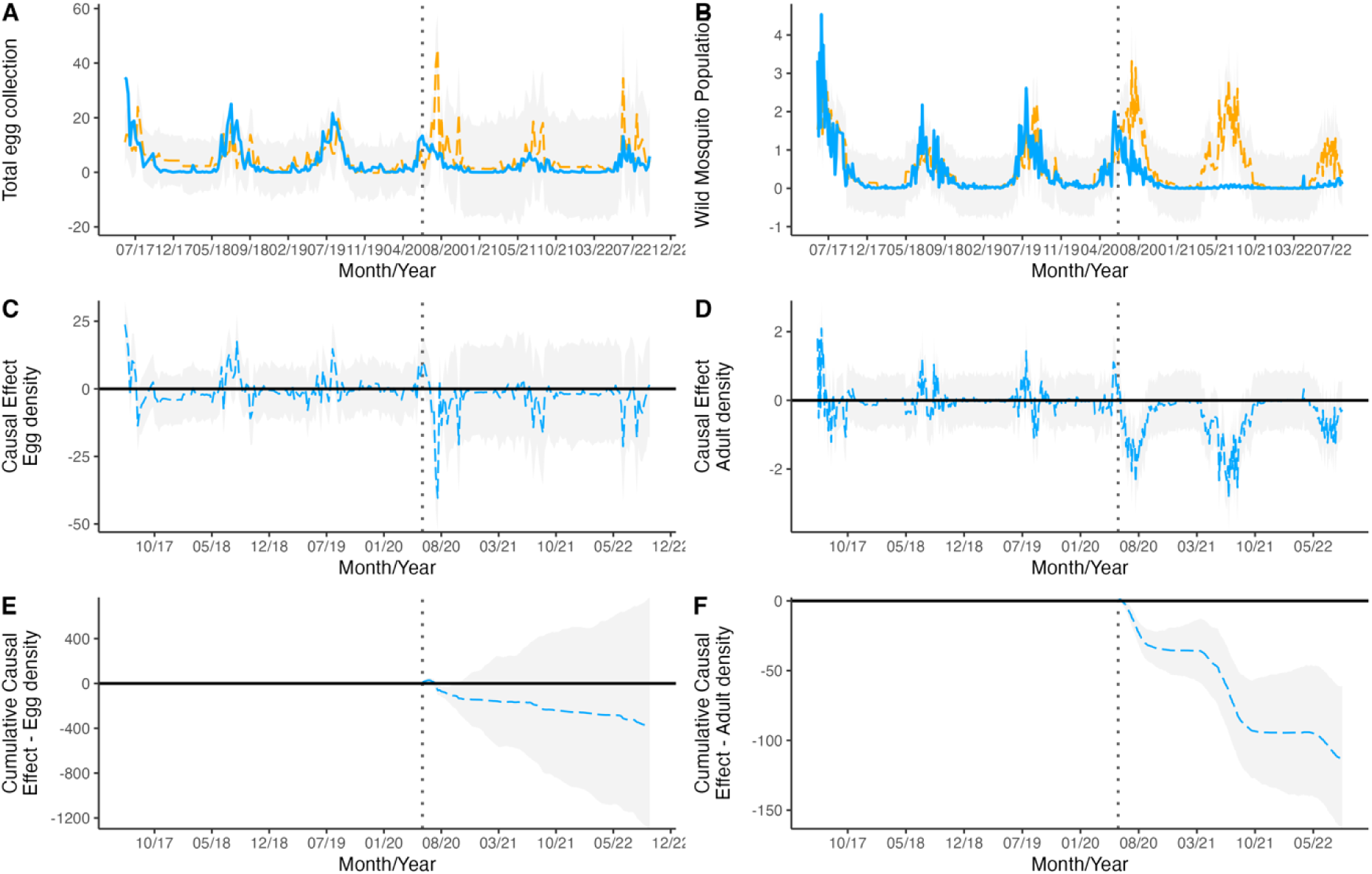
Complete causal impact effect analysis, with the comparison of the model and the observed data, followed by the model causal effect and the cumulative causal effect for the egg density (A, C, and E) and the adult density (B, D, and F) from phase 1 of the sterile male releases.

**S3 Figure.**
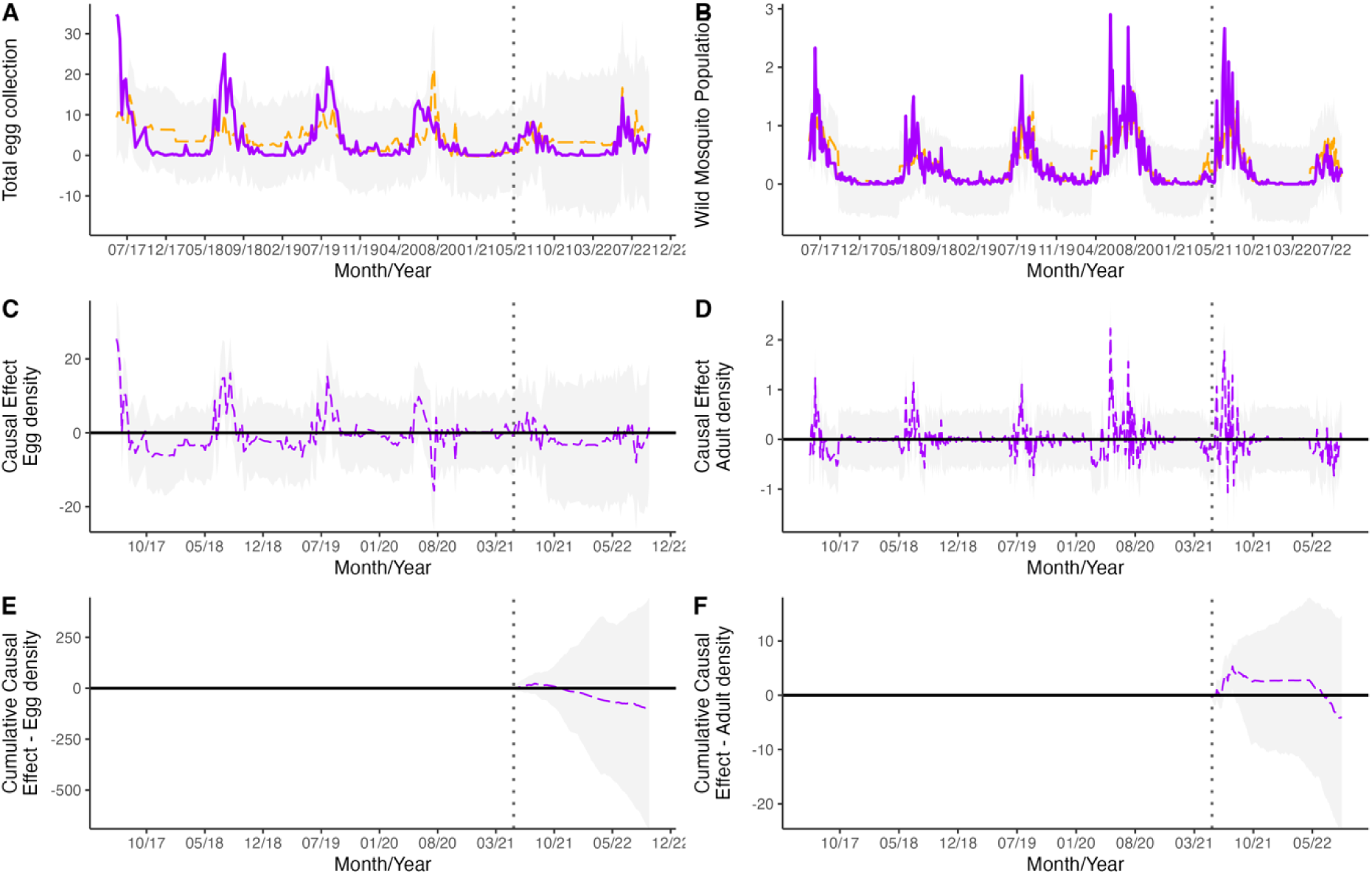
Complete causal impact effect analysis, with the comparison of the model and the observed data, followed by the model causal effect and the cumulative causal effect for the egg density (A, C, and E) and the adult density (B, D, and F) from phase 2 of the sterile male releases.

**S4 Figure.**
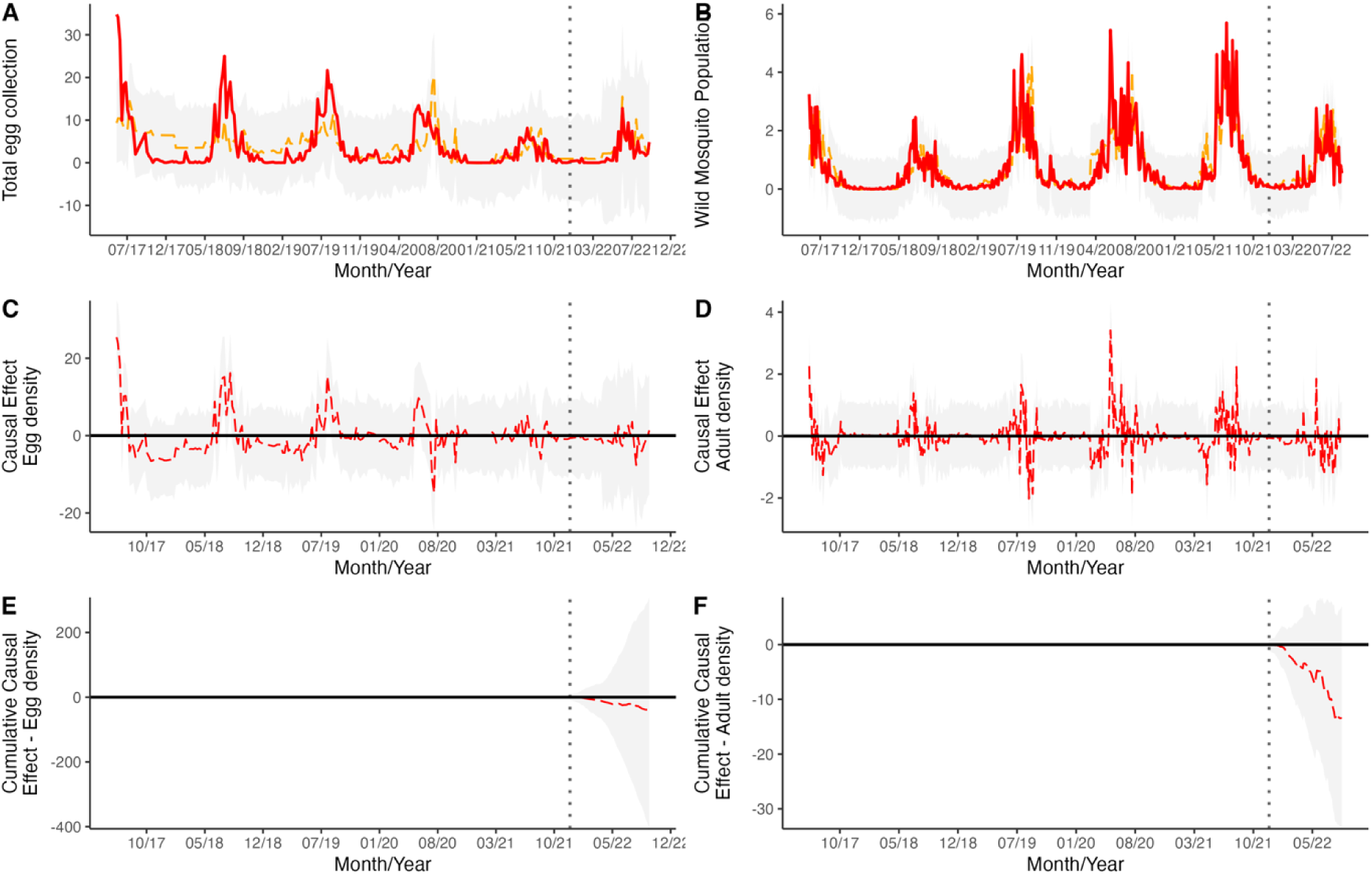
Complete causal impact effect analysis, with the comparison of the model and the observed data, followed by the model causal effect and the cumulative causal effect for the egg density (A, C, and E) and the adult density (B, D, and F) from phase 3 of the sterile male releases.

S5 Table. Statistical analysis output from the zero-inflated model with the emmeans.

S6 Text. Statistical analysis output from the Causal Impact Analysis (CIA).

S7 Table. Data on insecticide application during the releases in Captiva and Sanibel Islands.

S8 Text. Script for analysis in R.

## Notes

### Competing Interest Statement

The authors have declared no competing interest.

