## Supplementary material for "Suppression of *aedes aegypti* by sterile insect technique on captiva island, florida, usa from 2020 to 2022": S6 Text

CIA of Eggs Model Phase #1
Analysis report {CausalImpact}


During the post-intervention period, the response variable had an average value of approx. 2.43. In the absence of an intervention, we would have expected an average response of 5.65. The 95% interval of this counterfactual prediction is [-4.25, 13.70]. Subtracting this prediction from the observed response yields an estimate of the causal effect the intervention had on the response variable. This effect is -3.22 with a 95% interval of [-11.27, 6.68]. For a discussion of the significance of this effect, see below.

Summing up the individual data points during the post-intervention period (which can only sometimes be meaningfully interpreted), the response variable had an overall value of 279.00. Had the intervention not taken place, we would have expected a sum of 649.23. The 95% interval of this prediction is [-488.68, 1575.61].

The above results are given in terms of absolute numbers. In relative terms, the response variable showed a decrease of -80%. The 95% interval of this percentage is [-395%, +402%].

This means that, although it may look as though the intervention has exerted a negative effect on the response variable when considering the intervention period as a whole, this effect is not statistically significant, and so cannot be meaningfully interpreted. The apparent effect could be the result of random fluctuations that are unrelated to the intervention. This is often the case when the intervention period is very long and includes much of the time when the effect has already worn off. It can also be the case when the intervention period is too short to distinguish the signal from the noise. Finally, failing to find a significant effect can happen when there are not enough control variables or when these variables do not correlate well with the response variable during the learning period.

The probability of obtaining this effect by chance is p = 0.234. This means the effect may be spurious and would generally not be considered statistically significant. 


CIA of Adult Model Phase #1
Analysis report {CausalImpact}


During the post-intervention period, the response variable had an average value of approx. 0.13. By contrast, in the absence of an intervention, we would have expected an average response of 0.65. The 95% interval of this counterfactual prediction is [0.41, 0.88]. Subtracting this prediction from the observed response yields an estimate of the causal effect the intervention had on the response variable. This effect is -0.52 with a 95% interval of [-0.75, -0.28]. For a discussion of the significance of this effect, see below.

Summing up the individual data points during the post-intervention period (which can only sometimes be meaningfully interpreted), the response variable had an overall value of 29.25. By contrast, had the intervention not taken place, we would have expected a sum of 142.51. The 95% interval of this prediction is [90.31, 192.61].

The above results are given in terms of absolute numbers. In relative terms, the response variable showed a decrease of -79%. The 95% interval of this percentage is [-85%, -68%].

This means that the negative effect observed during the intervention period is statistically significant. If the experimenter had expected a positive effect, it is recommended to double-check whether anomalies in the control variables may have caused an overly optimistic expectation of what should have happened in the response variable in the absence of the intervention.

The probability of obtaining this effect by chance is very small (Bayesian one-sided tail-area probability p = 0.002). This means the causal effect can be considered statistically significant. 
NULL


Summary of Eggs Model Phase #2
Analysis report {CausalImpact}


During the post-intervention period, the response variable had an average value of approx. 2.53. In the absence of an intervention, we would have expected an average response of 3.95. The 95% interval of this counterfactual prediction is [-3.26, 12.63]. Subtracting this prediction from the observed response yields an estimate of the causal effect the intervention had on the response variable. This effect is -1.42 with a 95% interval of [-10.10, 5.79]. For a discussion of the significance of this effect, see below.

Summing up the individual data points during the post-intervention period (which can only sometimes be meaningfully interpreted), the response variable had an overall value of 172.20. Had the intervention not taken place, we would have expected a sum of 268.81. The 95% interval of this prediction is [-221.66, 859.06].

The above results are given in terms of absolute numbers. In relative terms, the response variable showed an increase of +24%. The 95% interval of this percentage is [-521%, +1356%].

This means that, although the intervention appears to have caused a positive effect, this effect is not statistically significant when considering the entire post-intervention period as a whole. Individual days or shorter stretches within the intervention period may of course still have had a significant effect, as indicated whenever the lower limit of the impact time series (lower plot) was above zero. The apparent effect could be the result of random fluctuations that are unrelated to the intervention. This is often the case when the intervention period is very long and includes much of the time when the effect has already worn off. It can also be the case when the intervention period is too short to distinguish the signal from the noise. Finally, failing to find a significant effect can happen when there are not enough control variables or when these variables do not correlate well with the response variable during the learning period.

The probability of obtaining this effect by chance is p = 0.363. This means the effect may be spurious and would generally not be considered statistically significant. 
NULL


Summary of Adult Model Phase # 2
Analysis report {CausalImpact}


During the post-intervention period, the response variable had an average value of approx. 0.29. In the absence of an intervention, we would have expected an average response of 0.32. The 95% interval of this counterfactual prediction is [0.17, 0.49]. Subtracting this prediction from the observed response yields an estimate of the causal effect the intervention had on the response variable. This effect is -0.033 with a 95% interval of [-0.20, 0.12]. For a discussion of the significance of this effect, see below.

Summing up the individual data points during the post-intervention period (which can only sometimes be meaningfully interpreted), the response variable had an overall value of 36.29. Had the intervention not taken place, we would have expected a sum of 40.41. The 95% interval of this prediction is [21.59, 60.88].

The above results are given in terms of absolute numbers. In relative terms, the response variable showed a decrease of -4%. The 95% interval of this percentage is [-40%, +68%].

This means that, although it may look as though the intervention has exerted a negative effect on the response variable when considering the intervention period as a whole, this effect is not statistically significant, and so cannot be meaningfully interpreted. The apparent effect could be the result of random fluctuations that are unrelated to the intervention. This is often the case when the intervention period is very long and includes much of the time when the effect has already worn off. It can also be the case when the intervention period is too short to distinguish the signal from the noise. Finally, failing to find a significant effect can happen when there are not enough control variables or when these variables do not correlate well with the response variable during the learning period.

The probability of obtaining this effect by chance is p = 0.332. This means the effect may be spurious and would generally not be considered statistically significant. 
Length  Class   Mode 
     0   NULL   NULL 


Summary of Eggs Model Phase #3
Analysis report {CausalImpact}


During the post-intervention period, the response variable had an average value of approx. 2.28. In the absence of an intervention, we would have expected an average response of 3.23. The 95% interval of this counterfactual prediction is [-5.29, 12.59]. Subtracting this prediction from the observed response yields an estimate of the causal effect the intervention had on the response variable. This effect is -0.96 with a 95% interval of [-10.32, 7.57]. For a discussion of the significance of this effect, see below.

Summing up the individual data points during the post-intervention period (which can only sometimes be meaningfully interpreted), the response variable had an overall value of 88.74. Had the intervention not taken place, we would have expected a sum of 126.12. The 95% interval of this prediction is [-206.40, 491.20].

The above results are given in terms of absolute numbers. In relative terms, the response variable showed a decrease of -80%. The 95% interval of this percentage is [-750%, +397%].

This means that, although it may look as though the intervention has exerted a negative effect on the response variable when considering the intervention period as a whole, this effect is not statistically significant, and so cannot be meaningfully interpreted. The apparent effect could be the result of random fluctuations that are unrelated to the intervention. This is often the case when the intervention period is very long and includes much of the time when the effect has already worn off. It can also be the case when the intervention period is too short to distinguish the signal from the noise. Finally, failing to find a significant effect can happen when there are not enough control variables or when these variables do not correlate well with the response variable during the learning period.

The probability of obtaining this effect by chance is p = 0.396. This means the effect may be spurious and would generally not be considered statistically significant. 
NULL


Summary of Adult Model Phase #3
Analysis report {CausalImpact}


During the post-intervention period, the response variable had an average value of approx. 0.64. In the absence of an intervention, we would have expected an average response of 0.84. The 95% interval of this counterfactual prediction is [0.54, 1.13]. Subtracting this prediction from the observed response yields an estimate of the causal effect the intervention had on the response variable. This effect is -0.19 with a 95% interval of [-0.49, 0.10]. For a discussion of the significance of this effect, see below.

Summing up the individual data points during the post-intervention period (which can only sometimes be meaningfully interpreted), the response variable had an overall value of 44.49. Had the intervention not taken place, we would have expected a sum of 57.93. The 95% interval of this prediction is [37.46, 78.15].

The above results are given in terms of absolute numbers. In relative terms, the response variable showed a decrease of -20%. The 95% interval of this percentage is [-43%, +19%].

This means that, although it may look as though the intervention has exerted a negative effect on the response variable when considering the intervention period as a whole, this effect is not statistically significant, and so cannot be meaningfully interpreted. The apparent effect could be the result of random fluctuations that are unrelated to the intervention. This is often the case when the intervention period is very long and includes much of the time when the effect has already worn off. It can also be the case when the intervention period is too short to distinguish the signal from the noise. Finally, failing to find a significant effect can happen when there are not enough control variables or when these variables do not correlate well with the response variable during the learning period.

The probability of obtaining this effect by chance is p = 0.1. This means the effect may be spurious and would generally not be considered statistically significant. 
Length  Class   Mode 
     0   NULL   NULL 
